# The inactive X chromosome accumulates widespread epigenetic variability with age

**DOI:** 10.1101/2023.03.10.532039

**Authors:** Yunfeng Liu, Lucy Sinke, Thomas H. Jonkman, Roderick C. Slieker, BIOS Consortium, Erik W. van Zwet, Lucia Daxinger, Bastiaan T. Heijmans

## Abstract

**Background:** Loss of epigenetic control is a hallmark of aging. Among the most prominent roles of epigenetic mechanisms is the inactivation of one of two copies of the X chromosome in females through DNA methylation. Hence, age-related disruption of X-chromosome inactivation (XCI) may contribute to the ageing process in women.

**Methods:** We analyzed 9,777 CpGs on the X chromosome in whole blood samples from 2343 females and 1688 males. We replicated findings in duplicate using one whole blood and one purified monocyte data set (in total, 991/924 females/males). We used double generalized linear models (DGLM) to detect age-related differentially methylated CpGs (aDMCs), whose mean methylation level differs with age, and age-related variable methylated CpGs (aVMCs), whose methylation level becomes more variable with age.

**Results:** In females, aDMCs were relatively uncommon (n=33) and preferentially occurred in regions known to escape XCI. In contrast, many CpGs (n=987) were found to display an increased variance with age (aVMCs). Of note, the replication rate of aVMCs was also high in purified monocytes (95%), indicating that their occurrence may be independent of cell composition. aVMCs accumulated in CpG islands and regions subject to XCI. Although few aVMCs were associated with X-linked genes in all females studied, an exploratory analysis suggested that such associations may be more common in old females. In males, aDMCs (n=316) were primarily driven by cell composition, while aVMCs replicated well (94%) but were infrequent (n=37).

**Conclusions:** Age-related DNA methylation differences at the inactive X chromosome are dominated by the accumulation of variability.

## Background

Epigenetic alterations are one of the five primary hallmarks of ageing [1]. A primary role for epigenetic mechanisms is the inactivation of one copy of the X chromosome in females to maintain dosage equivalence between females who carry two copies of the X chromosome, and males who carry a single copy [2]. Since the X chromosome harbors hundreds of protein-coding genes, many of which are implicated in disease including cancer [3] and neurological diseases [4, 5], age-related epigenetic changes at the inactivated X chromosome may be relevant for female aging.

For autosomes, it is well-established that age-related epigenetic differences occur at the level of DNA methylation. Although DNA methylation is tightly involved in the process of X-chromosome inactivation (XCI) [6], much less is known about age-related DNA methylation differences at chromosome X. Two recent studies addressed this question in whole blood samples and reported sets of CpG dinucleotides whose methylation level was associated with age separately for males and females [7, 8]. However, there was a striking lack of overlap between the results of both studies. Moreover, the previous studies focused on the occurrence of differences in mean methylation with age [7, 8] while there is increasing attention for the accumulation of variability in DNA methylation with age, a phenomenon that appears to be relatively independent of cell composition changes with age [9-12]. Finally, previous analyses did not consider the XCI status of regions harboring age-related DNA methylation differences. This is important because approximately 15% of genes on the inactive X consistently escape XCI and the escape status of an additional 10% of genes varies between tissues and individuals [13-15]. Hence, age-related epigenetic differences near such genes may be unrelated to XCI. These outstanding questions may be solved by analyzing larger sample numbers with robust statistical methods to test both differences in mean and variance followed by in-depth genomic annotation to relate findings to XCI.

Here, we report on the analysis of multiple discovery and replication cohorts totaling 3,334 female and 2,612 male blood samples with methylation data on 9,777 CpGs mapping to X chromosome. We detected age-related differentially methylated CpGs (aDMCs), whose mean methylation level differs with age, and age-related variable methylated CpGs (aVMCs), whose methylation level becomes more variable with age, while accounting for the impact of blood cell composition and inflation of test statistics. Systematic annotation, interpretation, and integration with transcriptomics data show that the inactive X is primarily affected by the accumulation of variance in DNA methylation with age, while the differences in mean are common at the active X chromosome but depend on changes in cell counts with age.

## Methods

### Discovery Cohorts

To discover age-related differentially methylation CpGs (aDMCs) and age-related variably methylated CpGs (aVMCs), genome-wide DNA methylation data were generated in whole blood samples within the Biobank-based Integrative Omics Studies (BIOS) Consortium, which comprises six Dutch biobanks: Cohort on Diabetes and Atherosclerosis Maastricht (CODAM) [16], LifeLines (LL) [17], Leiden Longevity Study (LLS) [18], Netherlands Twin Register (NTR) [19], Rotterdam Study (RS) [20], and the Prospective ALS Study Netherlands (PAN) [21]. Discovery data used in this study consists of 4031 (2343/1688 females/males) unrelated individuals for which DNA methylation data was available (**Additional File 1: Table S1)**. For 3131 (1794/1337 females/males) of these individuals also RNA-seq data were available. Data linkage of the two data types was verified from genotype data using *OmicsPrint* [22]. In addition, data on age, sex and technical batches were available for each cohort.

The generation of DNA methylation data has been described previously [23]. In brief, 500 ng of genomic DNA was bisulfite converted by the EZ DNA Methylation kit (Zymo Research, Irvine, CA, USA), and 4 μl of bisulfite-converted DNA was measured on the Illumina HumanMethylation450 array using the manufacturer’s protocol (Illumina, San Diego, CA, USA). Preprocessing and normalization of the data were done using DNAmArray workflow previously developed by our group (https://molepi.github.io/DNAmArray_workflow/). First, original IDAT files were imported into by R package *minfi* [24], followed by sample-level quality control (QC) was performed using *MethylAid* [25]. Filtering of probes was based on detection P value (P < 0.01), number of beads available (≤ 2), or zero values for signal intensity. Normalization was done using functional normalization as implemented in *minfi* [24], using five principal components extracted using the control probes for normalization. All samples or probes with more than 5% missing were excluded. In addition, probes with ambiguously mapping or cross-reactive were removed [26]. Finally, 9,777 X-chromosome CpGs and 4,031 samples were included in discovery set. Prior to analysis, we used *Combat* function of the SVA package to remove residual batch effects between cohorts, with biobank as batch and age, sex and known technical batches as covariates [27]. To exclude a negative impact of non-normal distributions and outliers on the validity of our results, DNA methylation data were transformed by rank-inverse normal (RIN) transformation for each cohort and females and males separately [28, 29].

Detailed information on the generation and processing of the RNA-seq data can be found in previous work [30]. In short, globin transcripts were removed from whole blood RNA using the Ambion GLOBINclear kit and subsequently processed for RNA-sequencing using the Illumina TruSeq version 2 library preparation kit. RNA libraries were paired-end sequenced using Illumina’s HiSeq 2000 platform with a read length of 2⍰×⍰50 bp, pooling 10 samples per lane. Reads which passed the chastity filter were extracted with CASAVA. Quality control was done in three steps: initial QC was performed using FastQC (v0.10.1), adaptor sequences were removed using Cutadapt, and read ends with insufficient quality were removed with Sickle. Reads were aligned to the human genome (hg19) using STAR (v2.3.0e). To avoid reference mapping bias, all GoNL SNPs (http://www.nlgenome.nl/) with MAF > 0.01 in the reference genome were masked with N. Read pairs with at most 8 mismatches, mapping to at most 5 positions, were used. Gene counts were calculated by summing the total number of reads aligning to a gene’s exons according to Ensembl, version 71. Samples for which less than 70% of all reads mapped to exons were removed.

For this study, we analyzed protein-coding genes mapping to the X chromosome. Raw counts of genes were transformed to log counts per million (CPM) values. After filtering out lowly expressed genes from the dataset (median CPM < 1), 512 out of 830 genes remained and were used for further analysis. Similar to DNA methylation data, RNA-seq data was transformed by rank-inverse normal (RIN) transformation for each cohort and females and males separately [28, 29].

### Replication cohorts

Two external datasets on DNA methylation data were used to replicate our results: a whole blood dataset originated from Sweden Population Health study by Johansson et al [31] and a purified monocytes dataset originated from Multi-Ethnic Study of Atherosclerosis by Reynolds et al [32]. These datasets here were referred as Johansson Blood and Reynolds Monocytes. Detailed information is shown in **Additional file 1: Table S1**.

For the Johansson Blood dataset, raw IDATA files were downloaded from the Gene Expression Omnibus (GEO) database (GSE87571). Also, information on age and sex for was available from GEO for this data set. The processing and normalization procedure of DNA methylation data is same as above. After processing, 9,853 X-chromosome CpGs were available For The Reynolds Monocytes dataset, DNA methylation array data was available from GEO (GSE56046) as quantile normalization method implemented in *lumi* package [33]. The GEO accession also included data on age and purity data of isolated monocytes. The sex of samples was predicted by *DNAmArray* package. After quality control, 9,861 X-chromosome CpGs remained. Both replication cohorts included all 9,777 X-chromosome CpGs measured in the discovery cohort.

### Detecting aDMCs and aVMCs on X-chromosome

To detect DNA methylation differences in both mean (aDMCs) and variance (aVMCs) with age, we applied double generalized linear model (DGLM). DGLM is a fully parametric method that first estimates mean effects by the linear model and then variance effects by the dispersion sub-model [34]. DGLM iterates between models until convergence. We used DGLM as implemented in the *dglm* R package (https://cran.r-project.org/web/packages/dglm/index.html) to identify aDMCs and aVMCs on chromosome X separately for females and males. The mean model part of DGLM was used to identify aDMCs, correcting for known covariates (age, cohort, cell counts and technical batches, namely sentrix position and sample plate) and unknown covariates include 5 latent factors estimated by *SVA* package [27]. Blood cell composition was estimated using IDOL method as implemented in the R package *minfi* [24] and resulted in predicted fractions for CD8T cells, CD4T cells, NK cells, B cells, monocytes and granulocytes. Of note, granulocytes were excluded from the model to exclude collinearity so that the effect of this cell type becomes included in the intercept.

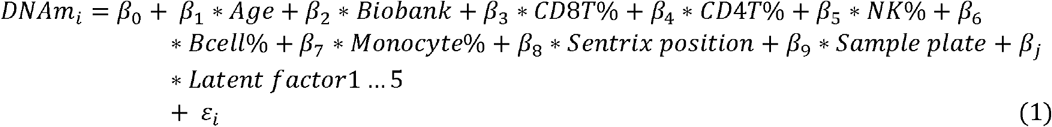

Where *DNAm*_*i*_ represent 9,777 X-chromosome CpGs methylation matrix, *Biobank* represent 6 cohorts comprising the data, *Sentrix position* represent sample position on the 450K array, *Sample plate* represent bisulfite plate and *β*_*j*_ represent regression coefficient for 5 latent factors estimated using *SVA* package [27], and *ε*_*i*_ represents the residual. Age-related differentially methylation changes were assessed by the parameter *β*_1_ in the mean model.

Then we estimated variance effect of age by the parameter *γ*_1_ in the dispersion sub-model:

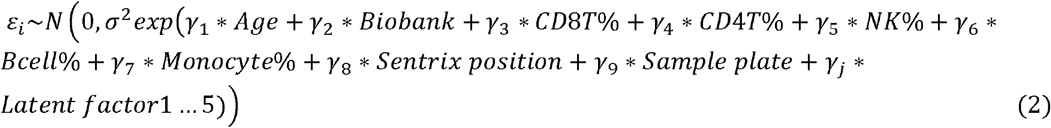

To reduce false positive findings, a Bayesian method implemented in R package *bacon* was used to correct the bias and inflation of test-statistics generated from mean model and dispersion sub-model of DGLM [35]. Statistically significance of the CpGs was determined by correcting multiple testing with a Bonferroni corrected P value <0.05.

To replicate findings, we repeated sex-stratified analysis with the same statistical model in Johansson Blood and Reynolds Monocytes datasets separately for all CpGs that were detected as aDMCs or aVMC in the discovery data set. For Johansson Blood data, cell counts were predicted the same way as for the discovery cohort. For the Reynolds Monocytes data, cell purity was included as covariate. CpGs were considered replicated if significant after correcting for multiple testing using the false discovery rate (FDR) (P_FDR_ < 0.05) [36] in both replication cohorts. The number of tests considered was the total number of aDMCs found in the discovery analysis (males + female) and likewise the total number of aVMCs.

Differentially methylated regions (DMRs) were called among replicated aDMCs and aVMCs in females and males separately by the *DMRfinder* algorithm [37] as implemented in the *DNAmarray* workflow (https://molepi.github.io/DNAmArray_workflow/). DMRs were defined as regions with at least 3 differentially methylated positions (DMPs) with an inter-CpG distance ≤ 1 kb, allowing maximum of three non-DMPs across a DMR [37].

### Genomic features of aDMCs and aVMCs

Replicated aDMCs and aVMCs were annotated according to 3 features. First, the CpGs involved were divided into three categories based on mean DNA methylation level: hypomethylated (β<0.25), intermediate methylated (0.25<β<0.7) and hypermethylated (β>0.7) since DNA methylation level is closely linked to XCI. Second, CpGs were mapped to 3 CpG island-related features: CpG islands (CGIs), CGI-shores and non-CGI regions, since XCI is associated with hypermethylation of CGIs [6]. The annotation was based on CGI-track downloaded from UCSC Genome Browser using version hg19 of the human genome. Shores were annotated as 2 kb regions flanking the CGIs and all remaining regions as non-CGI [37]. Finally, we annotated CpGs according to XCI status of the inactive X chromosome. Consensus XCI status calls per gene were previously defined [38] on the basis of three published studies [13, 14, 39] and resulted in genes classified into three categories: subject to XCI, escape XCI and variably escape XCI. These XCI status calls are referred to as meta-status calls [40]. For the annotation, we mapped CpGs to the nearest transcription start site (TSS) and when within 2kb from a TSS, the XCI status of the CpGs was set equal to the XCI status of the gene associated with the TSS. CpGs further away than 2kb from a TSS were not annotated because of uncertainty about their XCI status [41].

### Associations with X chromosome gene expression levels

To explore potential functional consequences of aDMCs and aVMCs, a linear regression model using the R package *limma* [42] was fitted to test for associations between the CpGs involved and the expression of 512 protein-coding genes mapping to the X chromosome. For this analysis, 3131 samples (1794/1337 females/males) from the BIOS consortium with both DNA methylation and gene expression were used. Known covariates included age, cohort, white blood cell composition as estimated with the R package *minfi* [24] and technical batches (eg, sentrix position, sample plate and flowcell number) were corrected in the linear regression model. Additionally, latent factors as estimated with the *SVA* package were included [27].

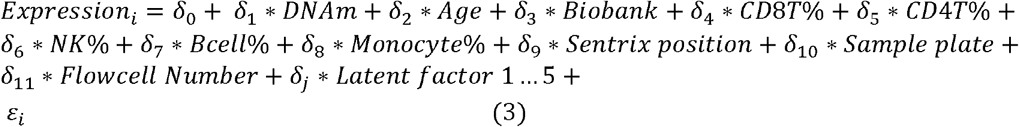

Where *Expression*_*i*_ represent 512 X-chromosome genes expression matrix, *Biobank* represent 6 cohorts comprising the data, *Flowcell Number* represent the HiSeq 2000 flowcell used for RNA-seq measurement, *δ* _*j*_ represent regression coefficient for 5 latent factors estimated by *SV A* package [27], and *ε*_*i*_ represents the residual. We used the R package *bacon* [35] to correct bias and inflation in the test statistics generated by this linear regression model and follow by multiple testing correction using the Bonferroni method (P_bonf_ < 0.05). Specifically, the bias and inflation of t-statistics was corrected for associations between each CpG and 512 investigated X-linked genes.

For X-linked genes associated with at least 1 CpG, we performed Gene Ontology enrichment analysis using the R package *clusterProfiler* using all other X-linked genes as background [43].

## Results

### Identification and replication of aDMCs and aVMCs on X-chromosome

To uncover age-related methylation changes on the X chromosome, we analyzed the methylation of 9,777 CpGs in whole blood using a discovery data set of 2343 females and 1688 males followed by the replication of findings both in an external whole blood dataset (388/341 females/males; **Table 1**) and an external dataset based on purified monocytes (603/583 females/males; **Table 1**). As expected, X-chromosome methylation levels were distinct for females and males (**Fig. 1a**). While the male X methylation pattern was similar to that of autosomes with the majority of CpGs being either hypo- or hypermethylated, the female X methylation pattern was trimodal with a significant proportion of CpGs showing intermediate DNA methylation levels. This stems from the fact that males carry a single active X, whereas females carry two copies of X, an active (Xa) and an inactive copy (Xi), where the Xi copy is mostly hypermethylated as part of the mechanisms ensuring XC I (**Fig. 1b**).

**Table 1.** Number of age-related Differentially Methylated CpGs (aDMCs) in the discovery and replication stage.

|  | Males |  |  | Females |  |  |
| --- | --- | --- | --- | --- | --- | --- |
|  | N | aDMCs (loss/gain) | Replication rate | N | aDMCs (loss/gain) | Replication rate |
| **Discovery** |  |  |  |  |  |  |
| BIOS Blood | 1688 | 1837 (718/1119) | - | 2343 | 80 (41/39) | - |
| **Replication** |  |  |  |  |  |  |
| Johansson Blood | 341 | 1371 (532/839) | 75% | 388 | 79 (41/38) | 99% |
| Reynolds Monocytes | 583 | 340 (195/145) | 25% | 603 | 34 (24/10) | 43% |
| Total | 2612 | 316 (175/141) | 17% | 3334 | 33 (24/9) | 41% |

**Figure 1.**
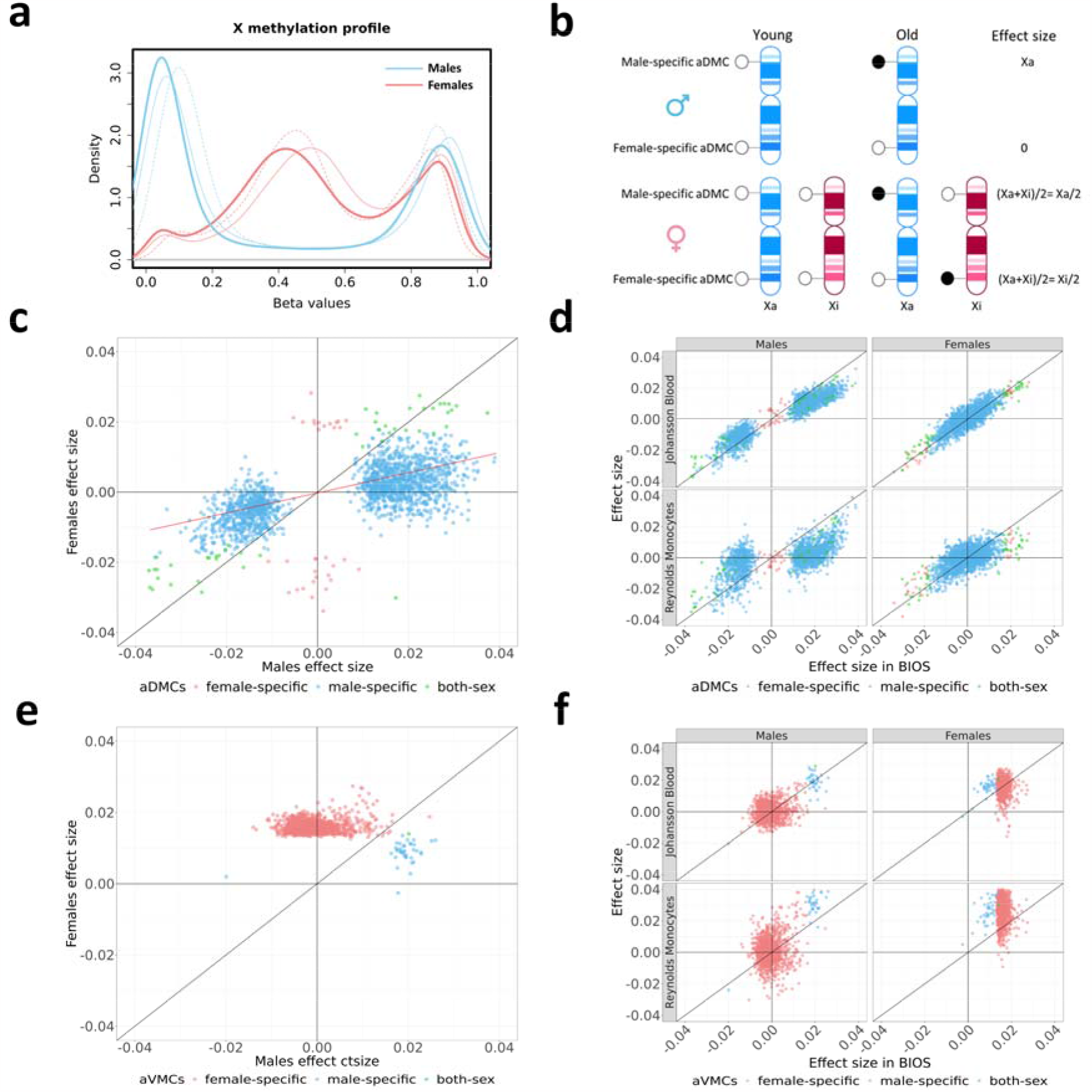
Identification and replication of aDMCs and aVMCs on the X chromosome. **a** Distribution of X-methylation in females and males based on discovery data (thicker solid line: BIOS Blood) and replication data (thinner solid line: Johansson Blood; thinner dash line: Reynolds Monocytes). The blue bimodal line and red trimodal line represent the distribution of X-methylation in males and females, respectively. **b** Schematic representation of how aDMCs specifically occurring on the Xa or Xi will affect the effect size as observed among males and females. Crucially, a study will have more statistical power to detect Xa-specific aDMCs in males than in females, because the presence of an addition Xa in females will dilute the effect size. c Scatter plot of effect sizes for aDMCs observed in males and females in the discovery data set. The Red regression line shows the relationship between the effect sizes in females for aDMC that are statistically significant in males only (**y=0.3x**) in line with a dilution of Xa-specific aDMCs in females due to the presence of a copy of Xi. Female-specific aDMCs, male-specific aDMCs and aDMCs statistically significant in both sexes are coloured by red, blue and green, respectively. **d** Scatter plot of aDMCs effect sizes in the discovery data sets and the Johansson Blood and Reynolds Monocytes replication datasets. **e** Scatter plot of effect sizes for aVMCs for females and males in the discovery data set. **f** Scatter plot of aVMCs effect sizes in the discovery data sets and the Johansson Blood and Reynolds Monocytes replication datasets. Female-specific aDMCs/aVMCs, male-specific aDMCs/aVMCs and aDMCs/aVMCs statistically significant in both sexes are coloured by red, blue and green, respectively. *Abbreviations: aDMCs* age-related differentially methylated CpGs, *aVMCs* age-related variable methylated CpGs

We first focused on CpGs whose mean methylation differed with age (aDMCs) using DGLM. T o confirm the robustness of this approach, we compared results with those obtained using a conventional model fitted using *limma* [ 40] and the effect sizes were virtually identical (**Additional file 1: Figure S1**). In the discovery data set, we identified 1837 aDMCs in males but only 80 in females (P_bonf_ < 0.05; **Table 1 and for a full list: Additional file 2: TableS2 and Table S3**). Of the aDMCs, 47 were shared between males and females. When inspecting the effect sizes for aDMCs, a striking pattern emerged (**Fig. 1c**): for female-specific aDMCs, the effect size in males was close to 0, whereas for male-specific aDMCs, the effect size in females was lower than in males, but correlated with that in males. This observation can be explained by the fact that males lack an Xi and, hence, female-specific aDMCs involving the Xi are fully absent in males (**Fig. 1c**). In contrast, male-specific aDMCs involve the Xa, and females also carry a Xa copy. Hence, the male-specific aDMCs are expected to be also present in females, albeit diluted by the presence of the unaffected Xi copy diluting the effect size two times (and therefore also increases the p-value). Indeed, we observed that the effect size for male-specific aDMCs was on average reduced by a factor of 0.3 in females (**Fig. 1c**).

To replicate our findings, we used two external datasets based of whole blood and purified monocytes samples (**Fig. 1d, Table 1**). We found that aDMCs effect sizes observed in discovery cohort were consistent with the effect size in Johannsson Blood dataset resulting in replication rates of 75% and 99% for male and female aDMCs, respectively. However, in the Reynolds data set based on purified monocytes, the replication rate of male aDMCs reduced to 25%, suggesting that the occurrence of aDMCs on Xa as observed in whole blood depends on age-related changes in blood cell composition that is not captured prediction of 6 cell types by the IDOL method. For females aDMCs, the replication rate was higher at 43%. Finally, 33 aDMCs in females and 316 aDMCs in males replicated in all 2 external data sets (**Table 1, Additional File1: Figure S2, Additional File2: Table S2 and Table S3**). Scatter plots for examples of replicated aDMCs are shown in **Additional file 1: Figure S3**. The examples included a female-specific aDMC (cg02991082) near a glucocorticoid-inducible molecule, *TSC22D3* (cg02991082), which is associated with anticancer immunosurveillance [44], and a male-specific aDMC cg13775533 near protocadherin 19, *PCDH19*, which is mainly expressed in the brain and linked to epilepsy (**Additional file 2: TableS2 and Table S3**) [45, 46]. Among the replicated male aDMCs, 17 DMRs were identified.

Next, we used our discovery data set to identify CpGs whose variability was associated with age (aVMCs) by fitting the dispersion sub-model of DGLM. In contrast to aDMCs, aVMCs were more common in females (n=1098) than males (n=39) with only one overlapping CpGs (P_bonf_ < 0.05, **Table 2, Additional File1: Table S4 and Table S5**). In line with the findings for aDMCs, male-specific aVMCs had an effect size in females that was approximately diluted by a factor 2 (mean effect sizes of 0.02 and 0.01 in males and females, respectively; **Fig. 1e**). Unlike aDMCs, aVMCs replicated surprisingly well in both the external whole blood and the external monocyte dataset for both sexes (>90%; **Table 2 and Fig. 1f**), which may be partly explained by earlier observations that aVMCs are less dependent on cell type composition of blood [9, 12, 47]. In total, 987 females aVMCs (located in 72 DMRs) and 37 males aVMCs replicated in both external data sets (**Table 2, Additional File1: Figure S4, Additional File2: Table S4 and Table S5**). Of note, all aVMCs were associated with an increase in variance with age except one, which decreased in variance with age specifically in males, also in the replication data sets. The CpG involved (cg25871420) and mapped to protocadherin 1 (*PTCHD1*) which associated with X-linked Intellectual Disability (**Additional file 1: Figure S3, Additional file 2: Table S4**) [48]. Examples of aVMCs increasing in variance with age in females included cg07876586 that mapped to Progesterone receptor membrane component 1 (*PGRMC1*), a gene associated with female-specific cancer types such as breast cancer [49] and ovarian cancer [50] (**Additional file 1: Figure S3, Additional file 2: Table S5**).

**Table 2.** Number of age-related Variably Methylated CpGs (aVMCs) in the discovery and replication stage.

|  | Males |  |  | Females |  |  |
| --- | --- | --- | --- | --- | --- | --- |
|  | N | aVMCs (loss/gain) | Replication rate | N | aVMCs (loss/gain) | Replication rate |
| **Discovery** |  |  |  |  |  |  |
| BIOS Blood | 1688 | 39 (1/38) | - | 2343 | 1098 (0/1098) | - |
| **Replication** |  |  |  |  |  |  |
| Johansson Blood | 341 | 37 (1/36) | 95% | 388 | 1034 (4/1030) | 94% |
| Reynolds Monocytes | 583 | 39 (1/38) | 100% | 603 | 1035 (0/1035) | 94% |
| Total | 2612 | 37 (1/36) | 95% | 3334 | 987 (0/987) | 90% |

### Annotation of X-linked aDMCs and aVMCs

The striking contrast in the occurrence of aDMCs and aVMCs in males and females indicated a differential involvement of Xa and Xi in the two types of age-related DNA methylation differences. A first feature associated with Xi is hypermethylation resulting in a predominance of intermediate DNA methylation levels in females (**Fig. 1a**). Compared to control CpGs, both aDMCs (P= 2×10^−6^) and aVMCs (P= 2×10^−16^) in females were preferentially intermediately methylated (**Fig. 2a**). Secondly, it is known that in particular CpG island (CGI) methylation is involved in XCI [6]. Nevertheless, aDMCs were depleted at CGIs in females and were even more common in non-CGI regions (**Fig. 2b**). However, aVMCs did preferentially occur at CGIs in females (P=4×10^−15^) while they were depleted in CGIs in males. Finally, we investigated previously reported XCI-status of Xi regions for aDMCs and aVMCs that were within 2kb of a TSS of genes with known XCI status to ensure validity of the XCI-status prediction (**Fig. 2c**). Annotated Female aDMCs all occurred outside regions subject to XCI (3 in escape and 2 in variably escape regions; P=2 ×10^−4^). In contrast, the large majority (83%) of annotated female aVMCs occurred in regions subject to XCI (P=3 ×10^−4^) in line with their enrichment at CGIs.

**Figure 2.**
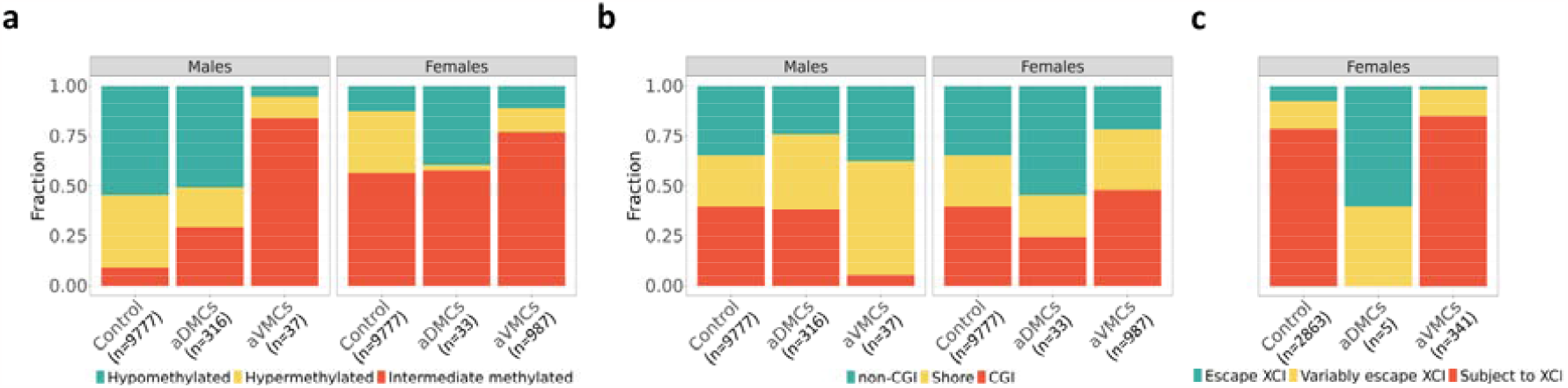
Annotation of aDMCs and aVMCs in males and females. **a** The mean methylation level of aDMCs and aVMCs in males and females (hypomethylated: β<0.25, intermediate methylated: 0.25<β<0.7, hypermethylated: β>0.7). b Fraction of aDMCs and aVMCs in CGI-related features (CGI, CGI-shore, and non-CGI). c XCI status annotation of aDMCs and aVMCs in females. Annotation was based on colocalization (<2kb) to TSS categorized as Subject to XCI, Escape XCI and Variably escape XCI. *Abbreviations: aDMCs* age-related differentially methylated CpGs, *aVMCs* age-related variable methylated CpGs, *CGI* CpG island, *XCI* X-chromosome inactivation

### Associations with X-linked gene expression

To explore whether the methylation level of aDMCs and aVMCs was associated with gene expression, we investigated the relationship between DNA methylation and 512 X-linked genes using a subset of 3131 individuals (1794/1337 females/males) from the discovery cohort for whom gene expression data were available in addition to DNA methylation data.

No associations with gene expression were observed for male aVMCs and female aDMCs. For 66 male-specific aDMCs, an association was observed with the expression of 19 X-lined genes (**Additional File3: Table S6**). The genes included several immune related genes such as *FRMPD3* and *CXCR3*. Other genes were disease-associated genes. For example, *DLG3* gene and *PIM2* gene were associated with X-linked intellectual developmental disorder [51] and Colorectal Adenocarcinoma [52] separately. For aVMCs, we only found that 3 female-specific CpGs associated with the expression of 2 X-linked genes (**Fig. 3a, Additional File3: Table S7**). The genes include *ALG13*, which is known to variably escape XCI [28] and *CDK16* which annotated as a gene escaping XCI. Neither aDMC-nor aVMC-associated genes were enriched for a certain biological pathway.

**Figure 3.**
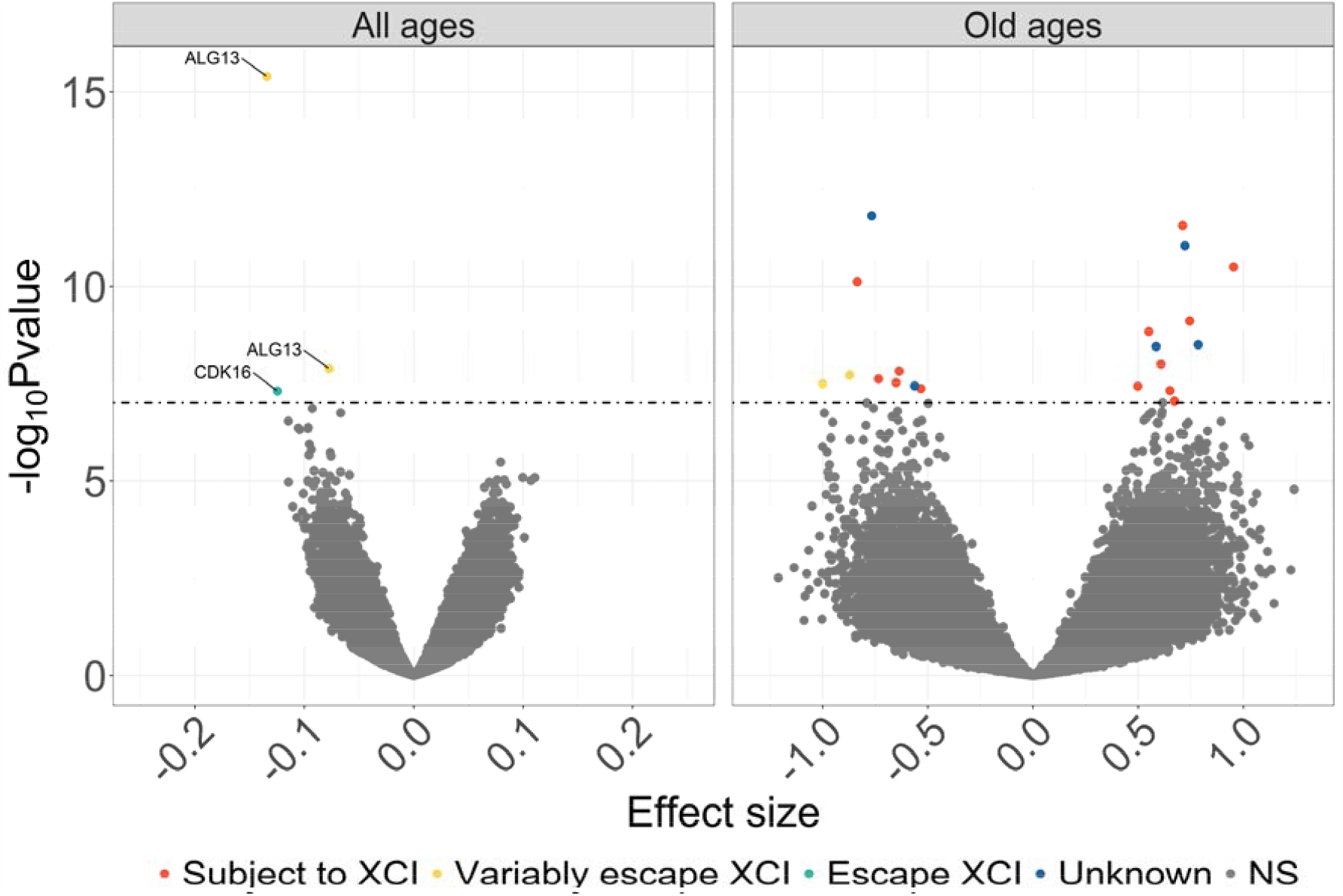
Volcano plot showing association between female-specific aVMCs methylation and X-chromosome gene expression in all females (n=1794, range 18-85 years) and in females older than 75 years (n=49). *Abbreviations*: *aVMCs* age-related variable methylated CpGs

One potential explanation for the small number of associations of aVMC CpGs with gene expression in females may be that they depend on more extreme DNA methylation levels that are only reached in old age, while our population had a mean age of x years (range, 18-85 years). To explore this hypothesis, we repeated our analysis in 49 elderly females (age≥75) out of the total of 1794 females with DNA methylation and gene expression data. In this very small subset, we found 19 female-specific aVMCs that were associated with the expression of 20 X-linked genes, 65% of which were subject to XCI (**Fig. 3b, Additional File3: Table S8**).

## Discussion

We report on a systematic analysis of age-related differences in X-chromosome methylation at the level of both differences in mean (age-related differentially methylated CpGs, aDMCs) and differences in variability (age-related variably methylated CpGs, aVMCs). We observed striking contrasts between the two types of age-related differences. aVMCs were common in females and rare in males and highly consistent across replication cohorts including in samples of purified monocytes. This suggests that the occurrence of X-linked aVMCs may be cell-intrinsic phenomenon in line with previous reports for autosomal aVMCs [9, 12, 47]. More commonly studied aDMCs, however, were rare in females and common in males, showed a poor replication rate, in particular in purified monocytes indicating that X-linked aDMCs were frequently driven by changes in blood cell composition with age. Further analysis supported the interpretation that aVMCs preferentially occur in regions subject to XCI on the inactive X, as characterized by enrichment at CGIs and the annotation of XCI status at aVMCs. Taken together, our data imply that DNA methylation marks involved in XCI commonly accumulate variability with age, hence suggesting that a gradual waning of epigenetic control at the inactive X may be a feature of female aging.

While aVMCs have previously not been reported for chromosome X, two previous studies reported on X-linked aDMCs. However, there was a great discrepancy between the aDMCs reported by both studies. First, Li et al. analyzed two discovery cohorts and found 559 and 1378 male-specific, and 1367 and 1148 female-specific aDMCs. Surprisingly, the overlap between findings of the two discovery cohorts was limited (34%-38%) and the replication rate in an external cohort was low (5-7%). We also found limited overlap between aDMCs from the two discovery cohorts and our replicated aDMCs (0.2%-3%; **Additional file 1: Figure S5**). The high frequency of aDMCs in females as compared with males in the two discovery cohorts of Li et al is at odds with the a priori expectation that the analysis of X methylation will have less power in females than males, because Xa-specific

DNA methylation differences will be diluted by the presence of an additional Xi and vice versa, while males carry a single Xa only (**Fig 1b**). Next, Kananen et al analyzed 5 studies and reported aDMCs that were observed in at least 2 out of 5 studies. Their findings overlapped to a higher degree with our replicated aDMCs (7%-17%, **Additional file 1:Figure S5**), and the overlap substantially increased when we applied a similar replication criterion to the Kananen aDMC set as in our current study, namely being a significant aDMC in 3 studies instead of 2 (30%-32%, **Additional file 1:Figure S5**). All in all, our aDMC results are more similar to that reported by Kananen et al [8] than that reported by Li et al [7]. Remaining differences between our study and the previous ones may be explained by differences in the study design and data analysis. Our total samples size (3334 females and 2612 males in discovery and replication cohorts) was substantially higher than that analyzed by Li (488 females and 488 males in discovery and replication cohorts) and Kananen (1191 females and 1240 males across 5 public datasets) [7, 8]. Moreover, among our replication cohort was a study based on purified monocytes instead of whole blood samples, rendering our results substantially less sensitive to blood cell type composition. Furthermore, we were more conservative in calling aDMCs. To correct for multiple testing, we applied Bonferroni correction in our discovery cohort and the false discovery rate for replication, whereas the previous studies used the more liberal false discovery rate for all analyses. In addition, we corrected for statistical inflation of test statistics (**Additional file 1: Figure S6**), a common problem in genomics studies which induces false positive findings [53]. Finally, we adhere to a strict replication scheme and report only those aDMCs that were replicated in all cohorts. It should be noted that in population studies like ours, findings in females are the average of Xa and Xi, which reduces the sensitivity to detect effects, but also the ability to definitely assign findings to either Xa or Xi. Nevertheless, our set of replicated aDMCs may be a robust starting point for further studies.

We observed that X-linked aVMCs are common in females and are associated with XCI. A main remaining question is whether aVMCs have functional relevance and how they can relate to the ageing process in females. We investigated the association of aVMC methylation with gene expression in females and only found two, *ALG13* and *CDK16*. This low yield may be explained in at least two ways. First, DNA methylation at aVMCs simply has little effect on gene expression. This would contrast with the assumed key role of DNA methylation in establishing and maintaining XCI. However, XCI involves multiple levels of epigenetic repression beyond DNA methylation (e.g. histone modifications). These other levels may not or be less affected with age. However, it could also be the case that age-related differences in DNA methylation levels may not reach a putative threshold that would result in loss of XCI. The latter links to a second potential explanation. It may be speculated that only when aVMCs reach extreme DNA methylation levels, for example in very old age, de-repression of regions normally subject to XCI occurs in a detectable way. Indeed, an exploratory analysis of a small subset of females aged 75 or over, suggested the presence of more associations of aVMCs with gene expression in old age. This observation requires replication in larger data sets. Such analyses will have to consider the fact that aVMCs result from stochastic phenomena whose occurrence in frequency and location may be highly individual-specific.

## Conclusions

Our analysis revealed that age-related DNA methylation changes at the inactive X chromosome are dominated by the accumulation of variability instead of commonly studied differences in mean. This implies that the epigenetic control of XCI may gradually wane with age. The putative functional impact of this phenomenon to female ageing may need to be studied in aged populations.

## Availability of data and materials

The BIOS whole blood samples is available from the European Genome-Phenome Archive (EGAC00001000277). Scripts for the analyses in this manuscript are available at GitHub [https://github.com/YunfengLUMC/Epigenetic_variability_chrX] under an open source MIT license. All external datasets used in this study are publicly available from Gene Expression Omnibus (GEO) under accession numbers GSE87571 [31] and GSE56046 [32].

## Acknowledgements

Samples were contributed by LifeLines, the Leiden Longevity Study, the Netherlands Twin Registry (NTR), the Rotterdam Study, the Genetic Research in Isolated Populations program, the Cohort on Diabetes and Atherosclerosis Maastricht (CODAM) study, and the Prospective ALS study Netherlands (PAN). We thank the participants of all aforementioned biobanks and acknowledge the contributions of the investigators to this study. We also thank Davy Cats and Hailiang Mei from the Sequencing Analysis Support Core (SASC) in the LUMC for their support at various stages of the project, including data pre-processing and maintenance of the Cloud based data analysis infrastructure. This work was carried out on the Dutch national e-infrastructure with the support of SURF Cooperative.

## Authors’ contributions

Conceptualization: BTH. Methodology: YL, BTH, LS, THJ, RCS. Formal analysis: YL. Resources: BIOS Consortium. Writing – original draft: YL, BTH. Writing – review and editing: BTH, LS, THJ, RCS, LD, EWvZ. Visualization: YL, BTH. Supervision: BTH. All authors have read and approved the final manuscript.

## Ethics declarations

### Ethics approval and consent to participate

The study was approved by the institutional review boards of the participating centers (CODAM, Medical Ethical Committee of the Maastricht University; LL, Ethics Committee of the University Medical Centre Groningen; LLS, Ethical Committee of the Leiden University Medical Center; PAN, Institutional Review Board of the University Medical Centre Utrecht; NTR, Central Ethics Committee on Research Involving Human Subjects of the VU University Medical Centre; RS, Institutional Review Board (Medical Ethics Committee) of the Erasmus Medical Center). All participants have given written informed consent, and the experimental methods comply with the Helsinki Declaration.

## Supplementary Information

*Additional File 1*.

Supplementary tables and figures. Contains **Fig. S1-S6** and **Table S1**.

*Additional File 2*.

Supplementary tables. Contains **Table S2-S5**.

Additional File 3.

Supplementary tables. Contains **Table S6-S8**.

