## Additional file 1 for "The inactive X chromosome accumulates widespread epigenetic variability with age"

**Table S1 Characteristics of cohorts used in present study.**

| Cohort | N | |  | Age (years) | | Accession |
| --- | --- | --- | --- | --- | --- | --- |
|  | Males | Females |  | Mean | Range |  |
| **Discovery**  **(BIOS Blood)** |  |  |  |  |  |  |
| CODAM | 86 | 74 |  | 65 | 50-79 | EGAC00001000277 |
| LL | 313 | 427 |  | 46 | 18-81 |  |
| LLS | 344 | 375 |  | 58 | 30-79 |  |
| NTR | 485 | 933 |  | 37 | 18-79 |  |
| PAN | 107 | 70 |  | 62 | 37-87 |  |
| RS | 353 | 464 |  | 68 | 38-87 |  |
| **Replication** |  |  |  |  |  |  |
| Johansson Blood | 341 | 388 |  | 47 | 14-94 | GSE87571 |
| Reynolds Monocytes | 583 | 603 |  | 60 | 44-83 | GSE56046 |

**Figure S1 Comparison of aDMCs effect size in both sex between DGLM and limma.**

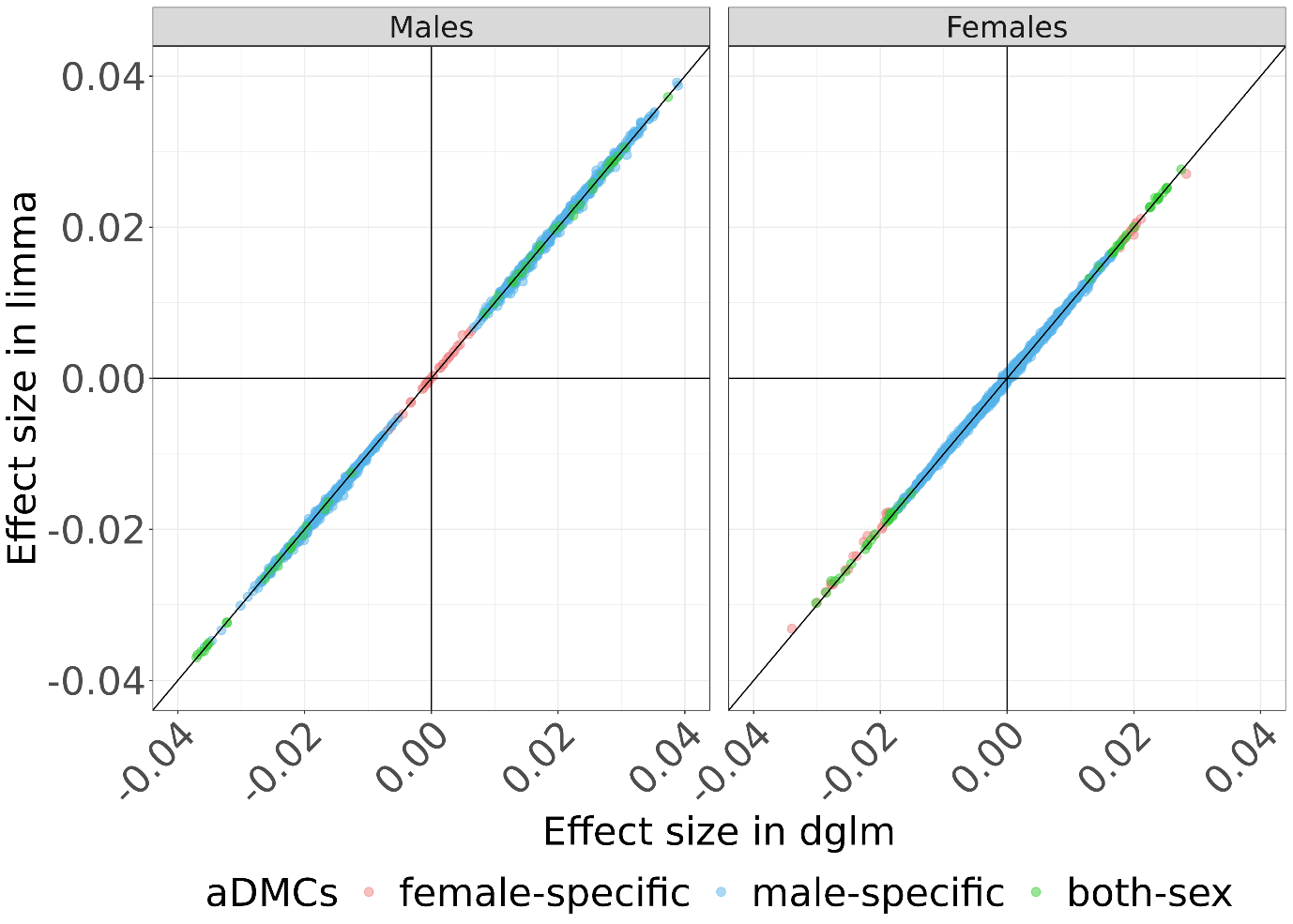

**Figure S2 Number of statistically significant overlapping aDMCs in males (a) and females (b) between discovery cohort (BIOS blood) and replication cohort (Johansson Blood and Reynolds Monocytes).** Orange column represent number of final validated aDMCs. *Abbreviations*: *aDMCs* age-related differentially methylated CpGs

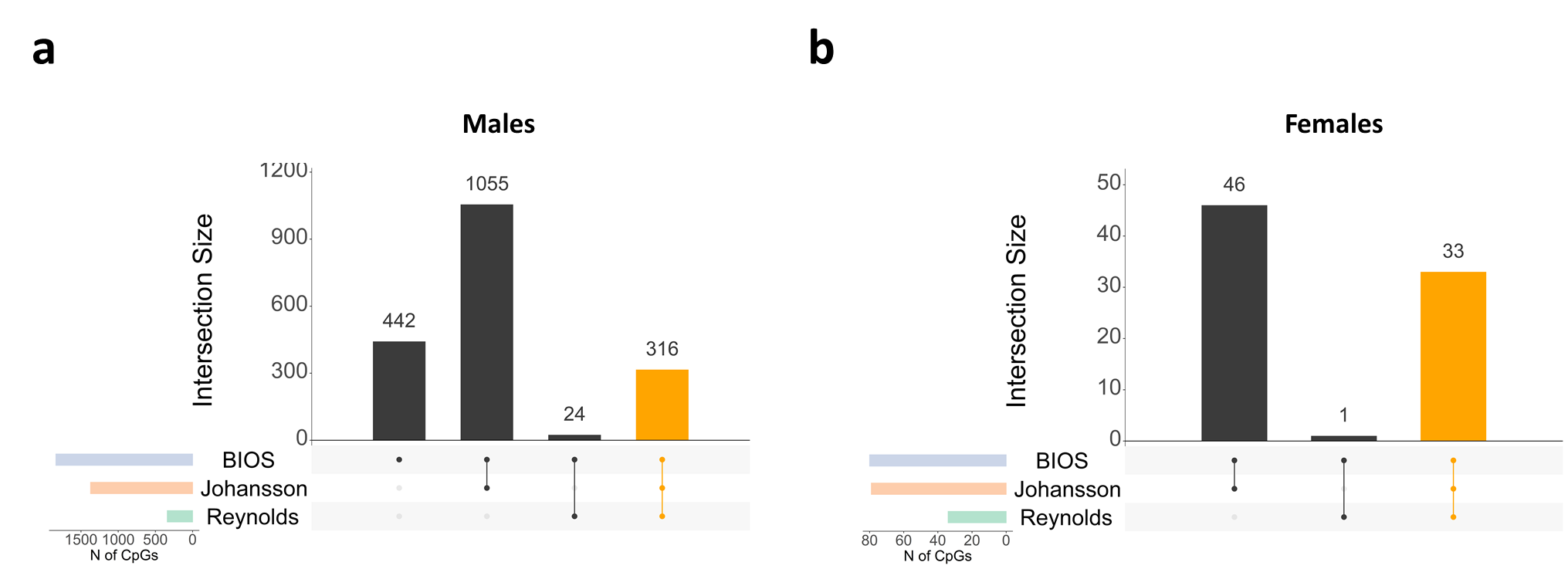

**Figure S3 Examples of replicated aDMCs and aVMCs in females and males.** Left scatter plot showing aDMCs that change in average DNA methylation with age. The aDMCs methylation (mean effects: middle line indicated) increased or decreased with age. Right scatter plot showing aVMCs methylation that change in variance with age. The aVMCs methylation variance (dispersion effect: extra two lines indicated) increased or decreased with age. DNA methylation value were rank-inverse normal transformed (y-axis). *Abbreviations*: *aDMCs* age-related differentially methylated CpGs, *aVMCs* age-related variable methylated CpGs

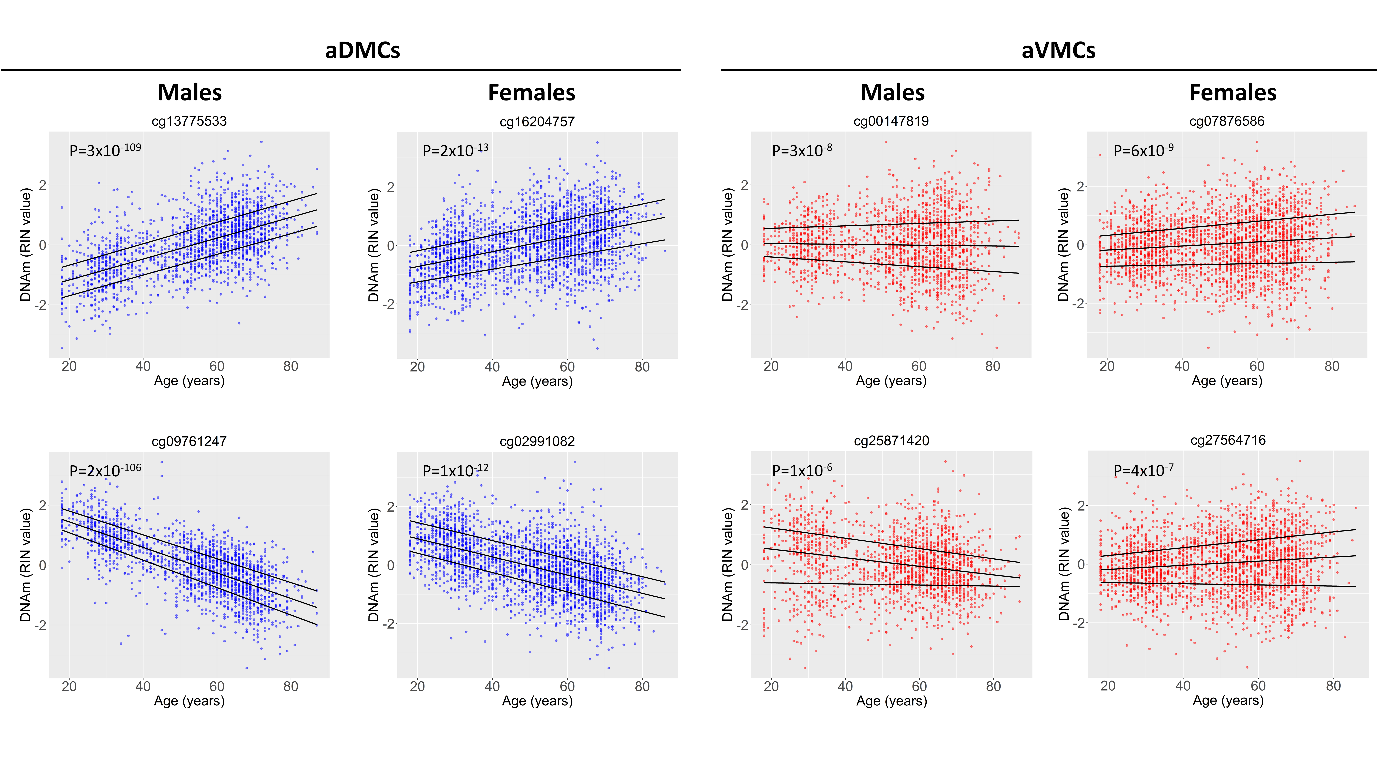

**Figure S4 Number of statistically significant overlapping aVMCs in males (a) and females (b) between discovery cohort (BIOS blood) and replication cohort (Johansson Blood and Reynolds Monocytes).** Orange column represent number of final validated aVMCs. *Abbreviations*: *aVMCs* age-related variable methylated CpGs

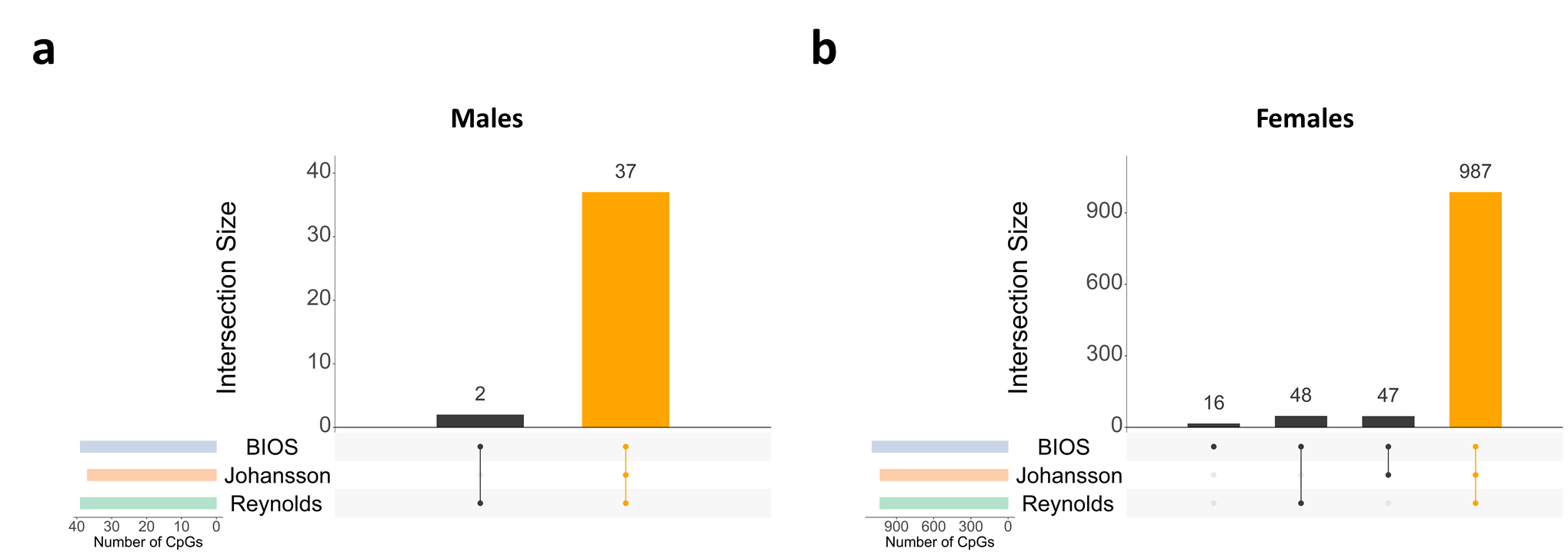

**Figure S5 Comparison of replicated aDMCs catalogue in our study with previous study.** a *Li* et al. *vs* *Kananen* et al. b *Li* et al *vs* Our study. c *Kananen* et al *vs* Our study.

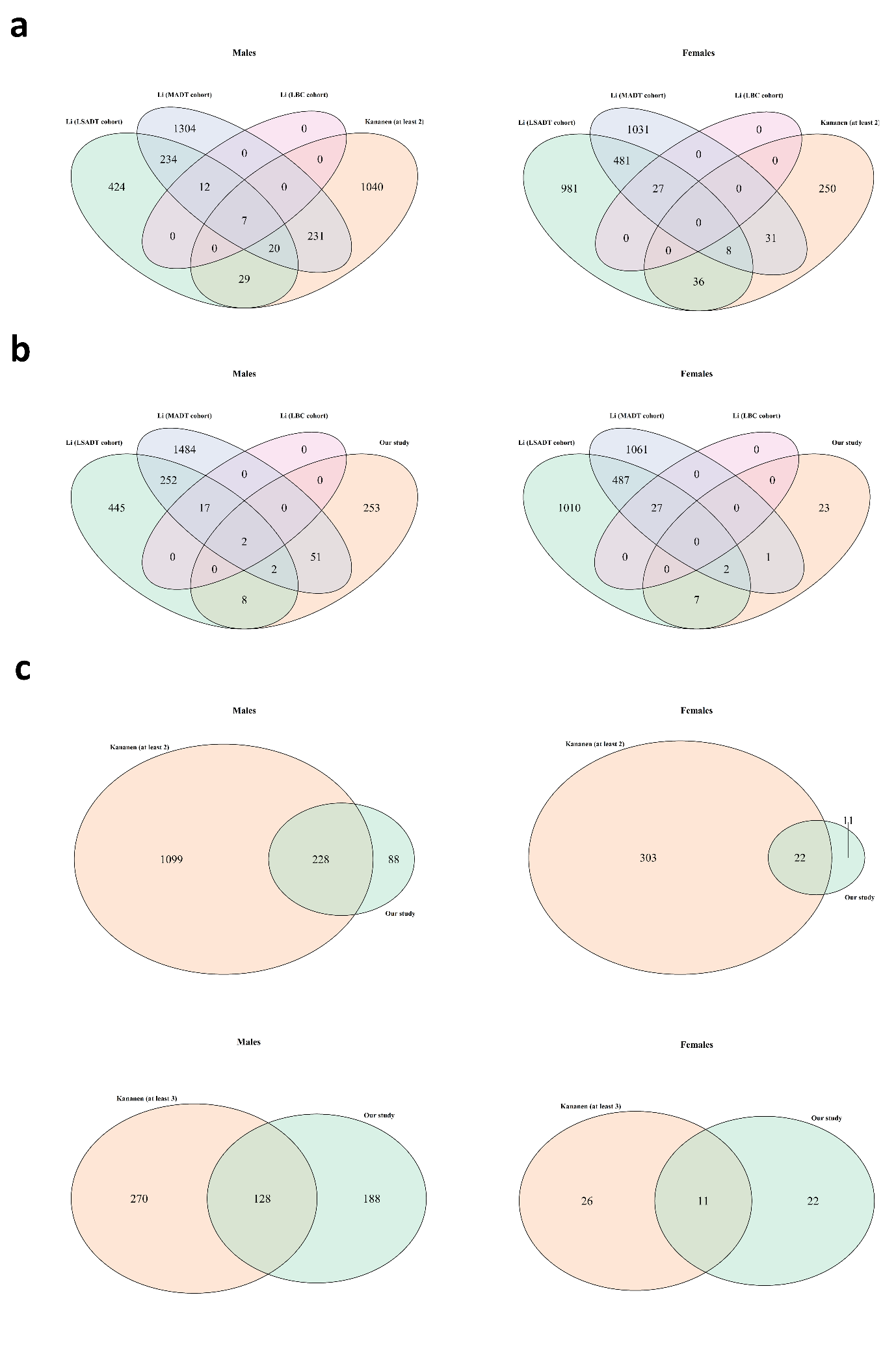

**Figure S6 Histogram of test statistics for aDMCs in males (a) and females (b) separately.** The black line represents the overall fit, the red line is the fit of empirical null distribution with estimated mean and variance. The green and blue lines represent the proportion of true positively and true negatively associations.

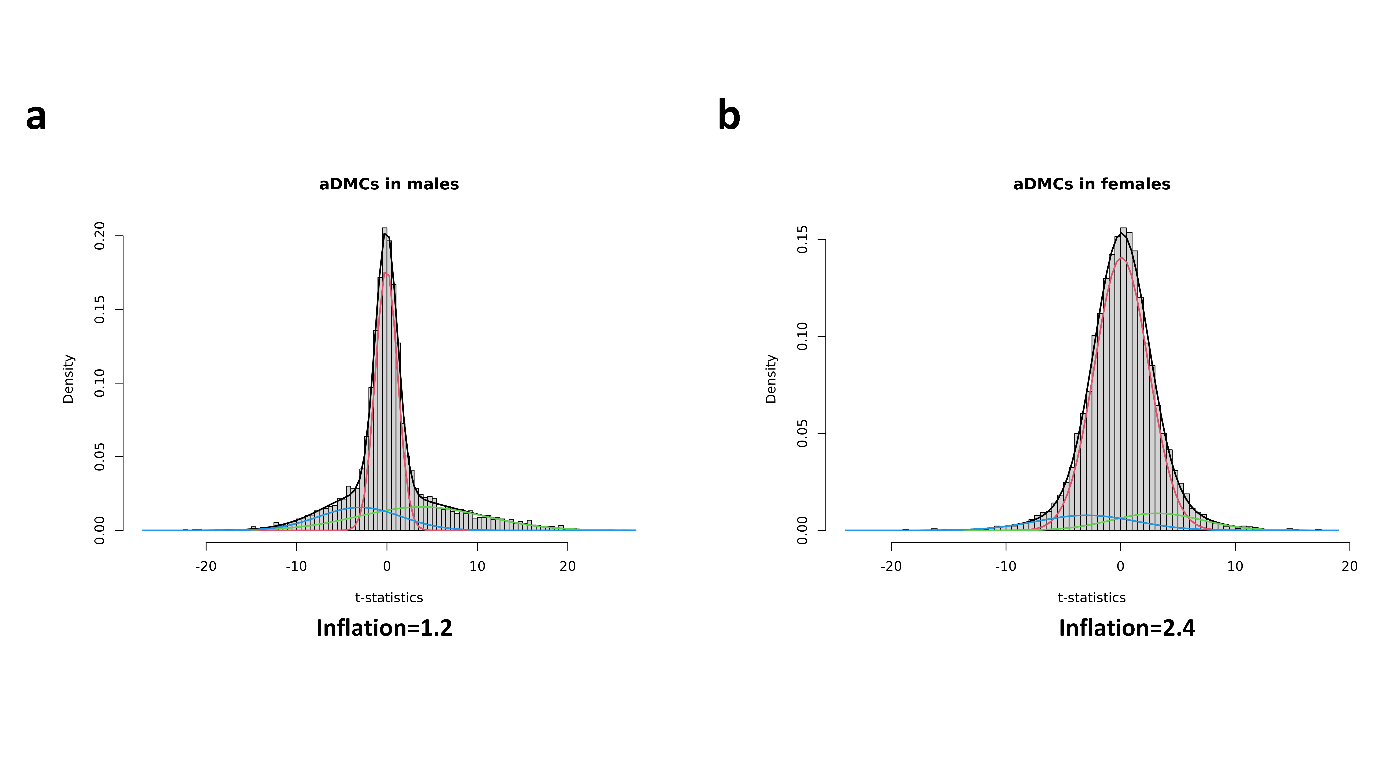
